# Unique allometry of group size and collective brain mass in humans and primates relative to other mammals

**DOI:** 10.1101/829366

**Authors:** Marcus J. Hamilton, Robert S. Walker

## Abstract

Group living is common in mammals, particularly in primates and humans. Across species, groups are social networks where co-residing members exchange information and balance trade-offs between competition and cooperation for space, resources, and reproductive opportunities. From a macroecological perspective, species-specific group sizes are ultimately constrained by body size, population density, and the environmental supply rate of home ranges. Here, we derive an allometric null model for group size in mammals based on individual energy demands and ecological constraints. Using Bayesian phylogenetic mixed models we show that primates exhibit unique allometries relative to other mammals. Moreover, as large-bodied primates, human hunter-gatherers have among the largest social groups of any mammal. We then explore the consequences of this unique social allometry by considering how mammalian brain size scales up in social groups that differ in size across mammals. We show similarly unique allometries in what we term the collective brain mass of social groups in primates relative to all other mammals. These results show that for a given body size primates have both larger brains and larger social networks than other mammals. Consequently, proportionally larger primate brains interact in proportionally larger social networks with important consequences for group cognition. We suggest that the size, scale, and complexity of human social networks in the 21^st^ century have deep evolutionary roots in primate ecology and mammalian brain allometry.

## 1. Introduction

Two of the most conspicuous features of the human species are large brains and intense sociality. How the two interact to influence cognition has become the focus of research across many disciplines (Clark and Chalmers, 1998; Dunbar, 1998; Richerson and Boyd, 2005; Dunbar and Shultz, 2007; Krubitzer, 2009; Woolley et al., 2010; Whiten and Erdal, 2012; Hutchins, 2014; Dennett, 2017; Everett, 2017; Graziano, 2017; Muthukrishna et al., 2018). Human brains are large, complex, and metabolically expensive, constituting ∼25% of the basal metabolic budget but only ∼2% of the body size. However, the computational returns on metabolic investment have been considerable. The initial doubling of hominin brain size to ∼800 cm^3^ in *Homo erectus* (*senso lato*) ∼2 MYA correlated with the expansion of the geographic range throughout Africa and southern Eurasia. The next major increase in brain size to ∼1,350 cm^3^ ∼300 KYA saw another expansion where modern humans replaced other hominins wherever they existed, eventually extending the human geographic range to include the majority of the planet’s terrestrial habitats. Humans began to genetically reengineer the biosphere ∼12 KYA by redirecting flows of environmental net primary production to net agricultural production, and ∼0.2 KYA humans leveraged thermodynamic principles to engineer machines to convert heat into work using fossilized biomass (Smil, 2008, 2019). Currently, the human species numbers about 7.7 billion, most of whom are connected by global communication networks, and now, through various technologies, have near-instant access to the majority of cultural knowledge accumulated over the last several thousand years. Humans now actively explore the solar system, are capable of manipulating matter at the smallest scales and gathering information at the largest. These innovations were facilitated not only by an initial increase in brain size and function, but by the intensity of social interactions.

The story of this evolutionary sequence is told, however incompletely, by the paleoanthropological, archaeological, and historical records (Christian, 2011). Explaining how and why humans are capable of such innovations is less clear. While the human brain is large for a mammal of our body size (∼7 times the predicted size of a mammal, and ∼3 times that of a primate), human cognition is not just a function of brain size, but brain specialization, where increasing size facilitates increasing diversity of function (Fuster, 1999; Elston et al., 2006; Striedter, 2006; Passingham, 2008; Barton, 2012; Bullmore and Sporns, 2012; Herculano-Houzel, 2012, 2016; Passingham and Wise, 2012; Roth and Dicke, 2012; Schneider, 2014; Wittmann et al., 2018). While the number of neurons in the mammalian brain increases with size, in primates an increasing proportion are located in the cerebral cortex (Herculano-Houzel, 2012). As the human brain is largely a linearly scaled-up primate brain, as large-bodied primates, humans have particularly neuron-rich cerebral cortices compared to other mammals (Herculano-Houzel, 2009, 2012). Consequently, in contrast to other primates, humans can mentalize at high orders, engaging in both self-reflection and on the mental state of others (Frith and Frith, 2006; Passingham, 2008; Tomasello, 2009). Humans can place themselves in alternative circumstances, set goals, plan ahead and monitor progress, weigh alternatives, engage in mental time travel, use schema to simulate outcomes, or use theories to predict probabilities, communicate directly with each other using language, learn complex tasks quickly, and use past events to inform actions in the present or future (Fuster, 1999; Passingham, 2008; Graziano, 2013; Tomasello, 2014; Everett, 2017). However, the intelligence of an individual human brain is a necessary, but not sufficient condition for human innovation; adaptation, innovation, and creativity results not from individuals per se, but from networks of interacting individuals (Hutchins, 1995, 2014; Richerson and Boyd, 2005; Whiten and Erdal, 2012; Bettencourt, 2013).

In this paper we take a macroecological approach to understanding the allometry of group size and brain size across mammals by combining principles of metabolic ecology (Brown et al., 2004) with Bayesian phylogenetic mixed models (Bürkner, 2017). We first derive and test an allometric null model of group size across mammals. Next, we consider the allometry of brain size scaling in social groups across mammals; we term this quantity *collective brain mass*. We do not assume the size of a social group necessarily equates to the degree of social complexity (Silk, 2007); the size of the network is the number of nodes (i.e., individuals), whereas the complexity would be the statistics of the network structure (i.e., connectomics). Here, we use the size of a network to quantify the collective brain mass within a group to which an individual brain contributes and interacts. Clearly, the nature of interactions varies widely within mammalian social groups, from prairie dog warning vocalizations to elephant infrasonic rumbling to human story-telling traditions. However, by definition, social species co-reside in groups of conspecifics who benefit, in one way or another, from living with others who share similar cognitive abilities and who use information received from others, intentional or not, to modify their behavior.

## 2. Theory and results

### a) Scaling relations for group size and collective brain mass

#### i) Data

We compiled a large database of mammalian ecology from published sources, including body size, basal metabolic rate, group size, home range, population density, and brain size (see *Electronic Supplementary Material (ESM)* for details). To quantify the scaling behavior, we use ordinary least squares regression models and Bayesian phylogenetic mixed models (BPMM) (Bürkner, 2017). In the BPMMs the dependent variable is weighted by the variance-covariance matrix of evolutionary relationships between all species, thus controlling for phylogenetically-structured residuals. Intercepts and slopes of the dependent variable are allowed to vary using mammalian order as the random factor. This modeling technique allows us to isolate scaling behavior within different mammal orders while controlling for their phylogenetic history. All data and code used in this paper are available as *Electronic Supplementary Material* attached to this paper.

#### ii) Theoretical development

Allometric relations are captured by power functions that take the general form

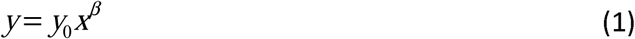

where *y* is a dependent variable, such as basal metabolic rate, brain size, group size, or home range size; *x* is an independent variable, commonly body size in allometry; *β* is the scaling exponent, an elasticity (*d* ln *y* / *d* ln *x*) capturing the proportional response of a change in *y* to a change in *x*; and *y*_0_ is a normalization constant. In biological systems when the independent variable is body size and the dependent variable is some measure of life history or physiology, *β* is commonly less than 1 (i.e., sublinear). As such, there is an inherent economy of scale in biological systems as mass-specific efficiency increases with size. In human social systems, properties relating to energy and infrastructure show similar economies of scale for similar reasons (Bettencourt et al., 2007; Hamilton et al., 2007a; Brown et al., 2011). However, when the independent variable is group size or population size and the dependent variable is a measure of collective productivity, such as wealth, innovation, crime, or incidence of disease, *β* is often greater than 1 (i.e., superlinear) (Bettencourt et al., 2007). This is because socioeconomic outputs are not the result of the number of people in a social network, but their interaction (Bettencourt, 2013); in a fully-connected unweighted social network the number of interactions, *c*, increases with network size, *n*, as *c* ∼ *n*^2^ and so connectivity increases multiplicatively with size. As such, social networks achieve increasing returns to scale from intensified rates of interaction as they grow in size. This paper combines these approaches to understand how collective social phenomena scale with body size across mammals.

### b) Basal metabolic rate

Group living begins with the metabolic energy required to support individual organisms. The basal metabolic rate is a fundamental rate in biology setting the energy demand of all biological functions (Brown et al., 2004). Across mammals, the relationship between the basal metabolic rate, *B*, and body size, *M*, is described by a power function *B* = *B*_0_*M* ^3/4^, where *B*_0_ is a mass-specific normalization constant; in our data the empirical scaling across all mammals from OLS is *B* ∝ *M* ^0.73^ (Figure 1A; Table 1). The BPMM shows no statistical difference in the metabolic scaling of primates to any other mammalian order (see Figure 1 A-C and *ESM*)). The human basal metabolic rate of ∼75 watts (the yellow star in figure 1A) is much as expected for a mammal of our body size (∼60,000 g, or 60 kg).

**Table 1.** Summary of OLS regression models and parameter estimates. For full details of ANOVA tables and results see the *Supplementary Information.*

| Figure | OLS Model | Constant | 95% CI | Slope | 95% CI | d.f. | P-value | r<sup>2</sup> |
| --- | --- | --- | --- | --- | --- | --- | --- | --- |
| 1A | $B \propto M$ | -3.94 | -3.88, -4.00 | 0.73 | 0.72, 0.74 | 793 | <0.001 | 0.95 |
| 1D | $W \propto M$ | -2.78 | -2.72, -2.84 | 0.73 | 0.72, 0.74 | 1366 | <0.001 | 0.96 |
| 1G | $W \propto B$ | -1.09 | 1.22, 1.14 | 0.98 | 0.96, 1.00 | 585 | <0.001 | 0.93 |
| 3A | $D \propto M$ | 8.63 | 8.35, 8.92 | -0.69 | -0.73, -0.65 | 1117 | <0.001 | 0.53 |
| 4A | $H \propto M$ | -9.82 | -10.19, -9.45 | 1.01 | 0.97, 1.06 | 950 | <0.001 | 0.64 |
| 4D | $H' \propto M$ | -9.32 | -9.78, -8.86 | 0.81 | 0.75, 0.87 | 785 | <0.001 | 0.49 |
| 5A | $N \propto M$ | -0.35 | -0.29, -0.51 | 0.17 | 0.15, 0.19 | 1205 | <0.001 | 0.21 |
| 6A | $G \propto M$ | -2.79 | -2.49, -3.09 | 0.87 | 0.85, 0.89 | 716 | <0.001 | 0.77 |

**Figure 1.**
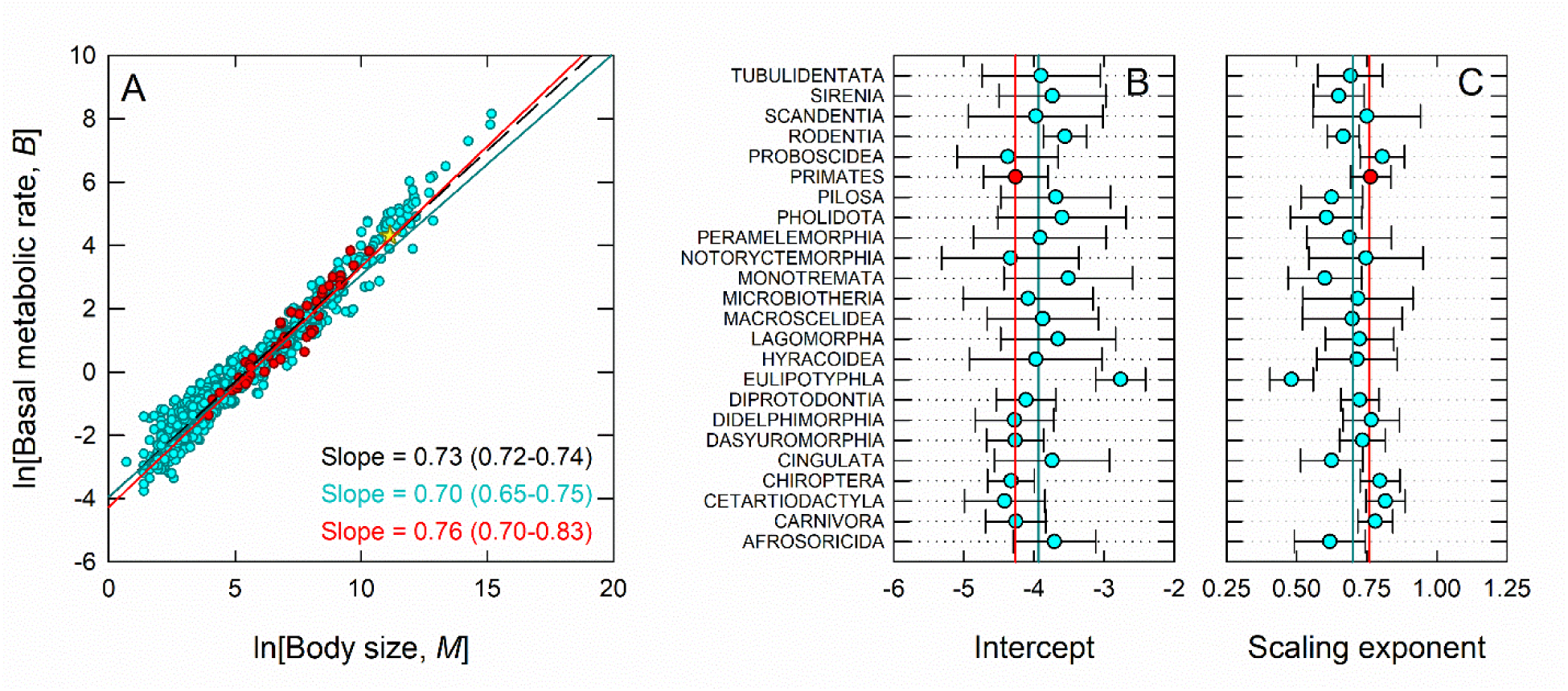
Basal metabolic rate and brain size allometries in mammals, primates and humans. Red data points, lines, and text are non-human primates, blue are non-primate mammals, and yellow diamonds are human. The red and blue lines are the scaling estimates from Bayesian phylogenetic mixed models and the dashed black line and text is the OLS model estimate of the slope. A) Scaling of basal metabolic rate in watts and body size in grams; B) Intercepts from the phylogenetic mixed model by order; C) Scaling exponents from the Bayesian phylogenetic mixed model by order.

### c) Group size, population density, and home range

At all body sizes some species live solitarily, but many live in social groups. Living in groups may help individuals maximize fitness by reducing predation risk, increasing foraging success, providing opportunities for alloparental care, increasing mating opportunities, or reducing mortality risk, for example (Krause et al., 2002; Silk, 2007; Beauchamp, 2013). On the other hand, group living may increase competition for space, resources, and mates, increase disease loads and the risk of free-riders. Ultimately the calculus of group size reduces to whether the net benefits of group living outweigh the costs (see Silk, 2007).

Human hunter-gatherer societies are organized into a complex hierarchy of social groups that form metapopulations (Figure 2) (Binford, 2001; Hamilton et al., 2007b; Bird et al., 2019). Interactions at all levels occur through fission-fusion dynamics that operate at different time-scales, from days to years (Hill et al., 2011, 2014). Using data from Binford (2001) we consider the five levels of hunter-gatherer social groups in Table 2.

**Table 2.** The five levels of hunter-gatherer social group sizes.

| Level | Group | Geometric mean | 95% CI | Sample |
| --- | --- | --- | --- | --- |
| 1 | Families | 4.48 | 4.31-4.67 | 116 |
| 2 | Dispersed bands | 15.60 | 14.68-16.58 | 228 |
| 3 | Aggregated bands | 53.66 | 49.86-58.29 | 298 |
| 4 | Regional populations | 165.32 | 152.25-181.00 | 214 |
| 5 | Ethnolinguistic metapopulations | 839.19 | 736.36-954.03 | 340 |

**Figure 2.**
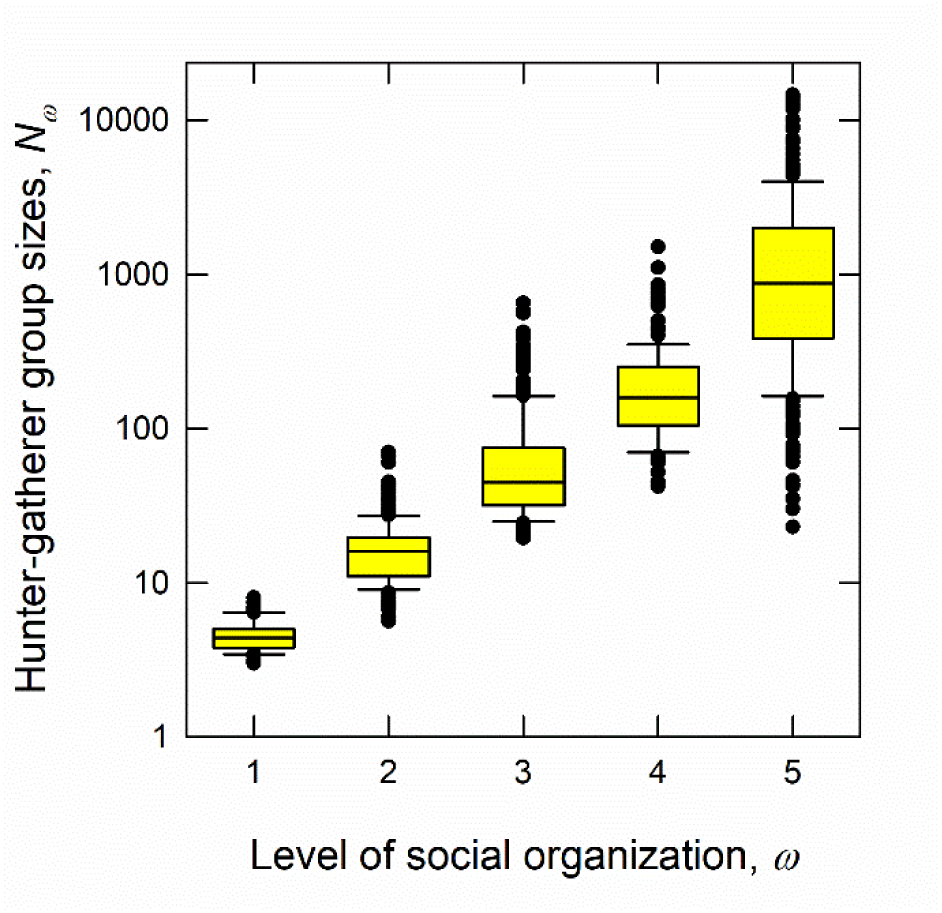
Boxplots of hunter-gatherer group sizes across five levels of social organization. The horizontal bars in the box bodies are medians, the height of the box is the middle 50% of the data, and the whiskers are +/-25%. At all scales of social organization hunter-gatherer group size are approximately normally distributed on the log scale, and so lognormally distributed on the linear scale. 1 = families, 2 = dispersed bands, 3 = aggregated bands, 4 = regional populations, and 5 = ethnolinguistic metapopulations (see Table 2 for details).

In mammals social group size, *N*, is defined as the number of individuals that co-reside within a shared home range, *H*, and is usually measured in the field as the average number of individuals with whom a set of focal individuals interact over a 24-hour time period (Jones et al., 2009). The allometry of group size is inherently noisy as there are solitary species at all body sizes and so there is no simple correlation between group size and body size that holds for all mammal species. In other words, while average group size may increase with body size, the variance will also increase as group size is bounded at *N* = 1 for all body sizes.

We derive a simple allometric null model for group size from the definitions of home range and population density. Let us assume the daily encounter rate of conspecifics, *λ*, is a function of the population density of conspecifics within a home range, and so the number of individuals encountered during a day, *N*, will be the product of the encounter rate, *λ*, and the home range size, *H*, both of which scale allometrically. Mammal population density, *D*, scales with body size as the inverse of metabolic rate; *D* = *N* / *A* ∝ *M* ^-3/4^, where *A* is a sampled area in km^2^. This allometric relationship is known as Damuth’s Law in ecology (Damuth, 1981). We see this scaling empirically in Figure 3A and Table 1. Note that primate population densities are shallower than the mammalian average (Figure 3C). In mammals, home range size scales linearly with body size; *H* ∝ *M* ^1^, (Peters, 1986; Kelt and Van Vuren, 2001; Jetz et al., 2004). We see this empirically in Figure 4A and Table 1. Figures 4B and C show the remarkable consistency of home range scaling across mammalian orders, where only Rodentia deviate from all other mammals. Combining these two results we have a null allometric expectation for group size, *N*:

**Figure 3.**
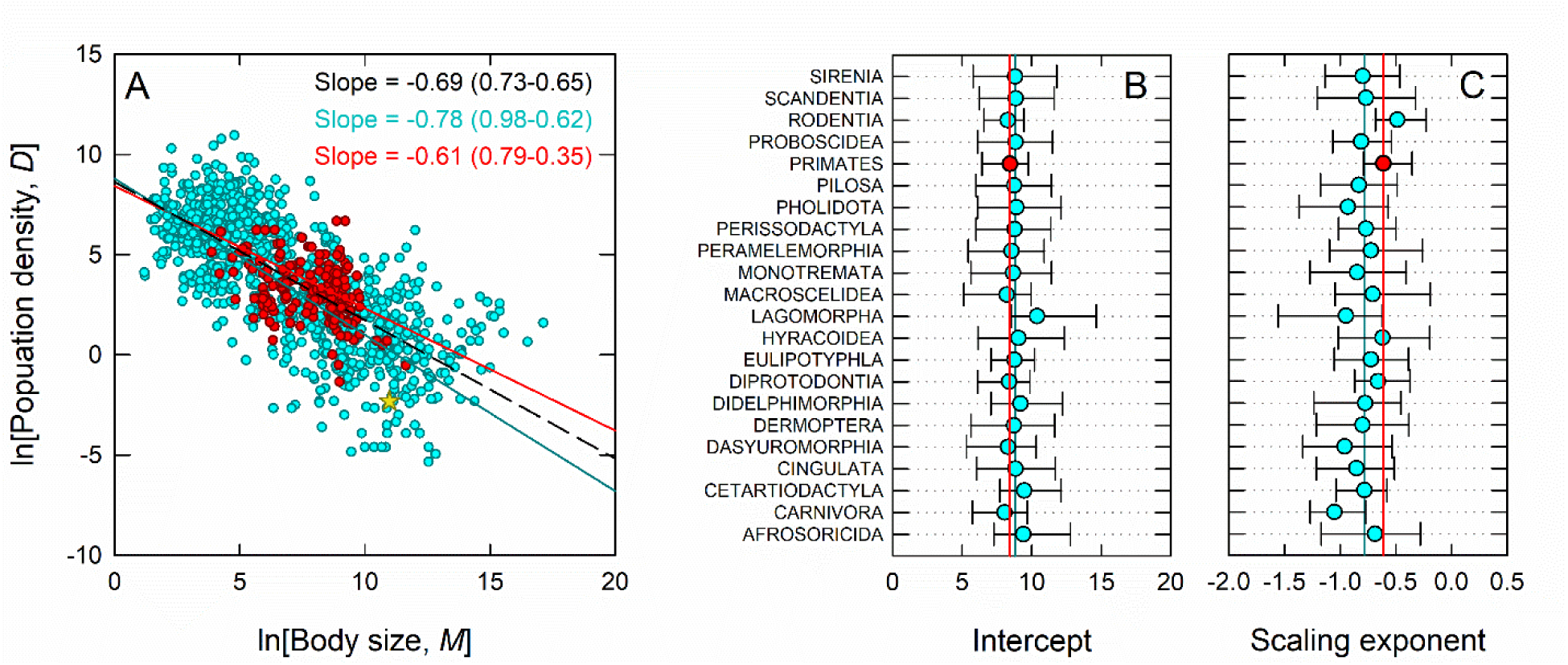
Population density allometries in mammals. A) The scaling of population density and body size; B) Intercepts for each order from the phylogenetic mixed model; and C) Slopes for each order.

**Figure 4.**
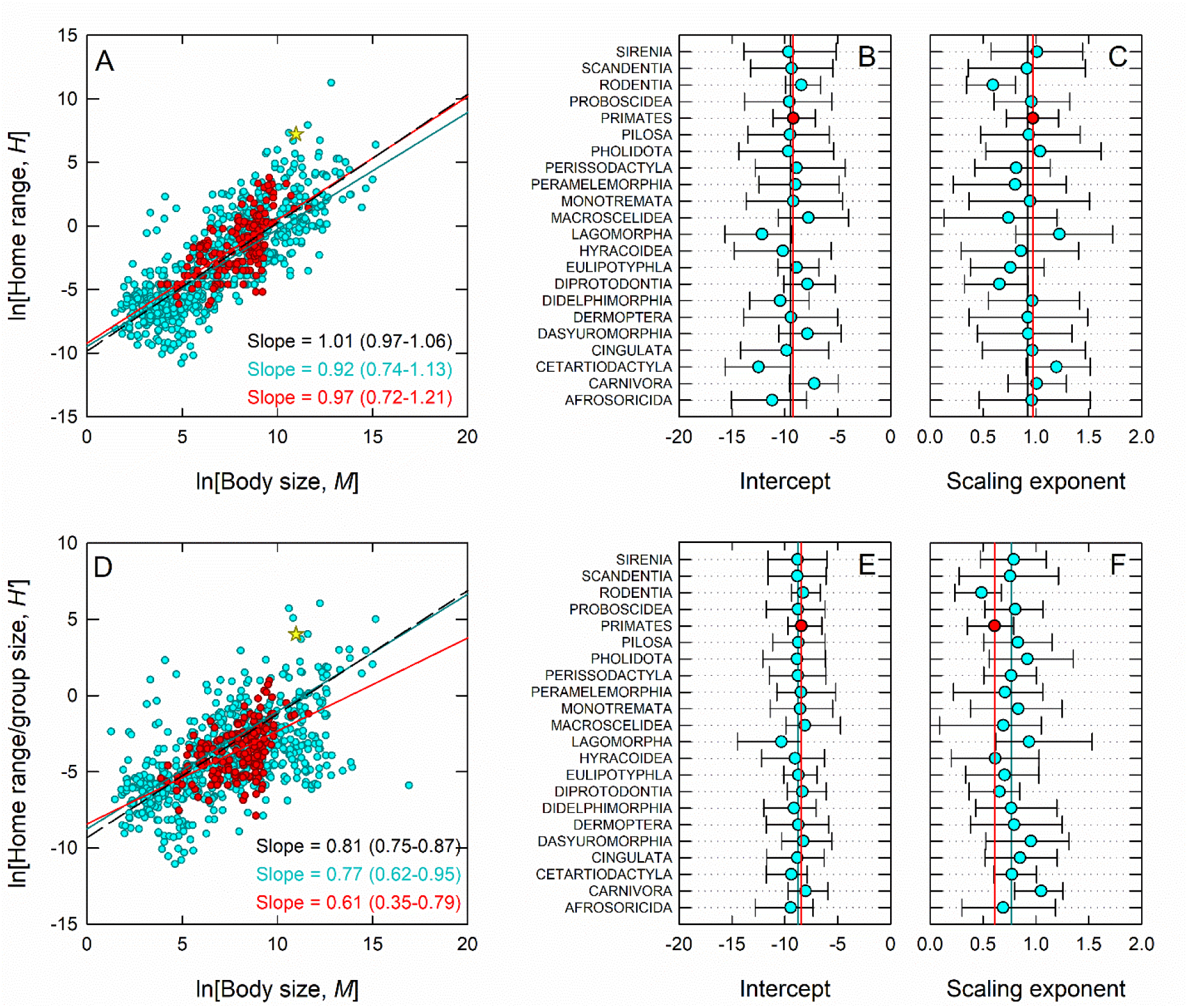
Home range allometries in mammals, primates and humans. A) The scaling of home range size and body size; B) Intercepts for each order from the phylogenetic mixed model; C) Slopes for each order; D-F) The scaling of individual home range size and body size, and phylogenetic mixed models results; G-I) the scaling of social group size and body size, and mixed effect model results.

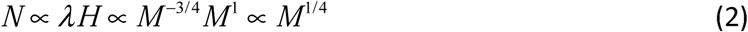

We first test this model using an approach developed by Jetz et al. (2004). The size of a home range, *H*, is the space required by a mammal to meet their metabolic demand, *B*, the resource supply rate from the local environment, *R*, and the spatial overlap with conspecifics; thus *H* = *B* / *α R*, where *α* is the proportion of the environmental resource supply rate used exclusively by an individual (Jetz et al., 2004). Therefore, the home range used by an individual in a group is estimated by dividing the home range, *H*, by group size, *N*. The group size-corrected home range is then defined as *H*′ = *H* / *N*. Following equation 2 and the allometry of home range the expected scaling of the group size-corrected home range is then

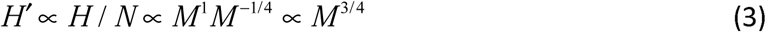

We find support for this predicted scaling in Figures 4D-F, where the scaling estimates are consistent this prediction. It is important to make two observation here. First, note that the group size-corrected home range, *H*′ = *H* / *N* (Figure 4D), is the inverse of population density, *D* = *N* / *A* (Figure 3A), and so Jetz et al.’s model directly links home range scaling with Damuth’s Law. Consequently, group size-corrected home ranges in primates deviate from the overall mammal scaling (Figures 4D and F) in the same way that we see in primate population densities (Figures 3A and C). Thus, for their body size primate species tend to be denser on the landscape than most other mammals because individuals overlap in space with conspecifics within their home ranges more so than other mammals. In other words, primate species in general sustain larger group sizes per unit area than other mammals with the exception of Rodentia and Eulipotyphla (shrews, moles, and hedgehogs) (Figure 4F). Interestingly, most species in the orders Primates, Rodentia, and Eulipotyphla are either arboreal or fossorial. As such, their home ranges include an additional vertical dimension that could explain the higher observed population densities.

In Figure 3A, the hunter-gatherer average population density (yellow star) was estimated from the geometric mean of total population sizes divided by the geometric mean of territory sizes using data from Binford (2001) (see ESM for data). Hunter-gatherers have relatively low population densities for mammals of their body size and have the lowest average population densities of any primate. In Figure 4A, hunter-gatherer home ranges are residential patch sizes estimated from mobility data in Binford (2001). The geometric mean distance of the average length of a residential move from the centroid of one patch to the centroid of an adjacent patch is *d* (see Hamilton et al., 2016). Assuming patch sizes are circular the patch size (i.e., home range), *H*, is estimated as *H* = *π* (*d* / 2)^2^. To estimate the group size-corrected home range we then divide this quantity by the average size of dispersed bands (level 2 in Table 2), as this is the number of individuals with which a hunter-gatherer will spend most of the year co-residing (Binford, 2001; Hill et al., 2011). Figures 4A and D show that hunter-gatherers have particularly large home ranges for mammals of their body size, and by far the largest of any primate (both absolutely and relatively).

Figure 5A shows that across mammals there is a noisy but statistically significant positive scaling relationship between group size and body size, *N* ∝ *M* ^0.17^ (Table 1), though shallower than the null expectation of *N* ∝ *M* ^1/4^. Figure 5C shows that primate group sizes increase with body size significantly faster than all other mammal orders, where *N* ∝ *M* ^0.39^, and significantly faster than the null prediction (the vertical black like in Figure 5C). As this scaling is steeper than the allometric null model, this means that in primate species individuals form groups that are larger than simply the average density of individuals encountered in their home range over a 24 hour period. In other words, this is strongly indicative of prosocial behavior as there must be additional behaviors that aggregate individuals into cooperative groups than simply shared space and random encounters. On the other hand, in all other mammalian orders, the scaling slope is less than 1/4 and so individuals form social groups at rates slower than the null model. Group formation is much more limited and constrained in most other mammalian orders than in primates. Figure 5A shows that largest group sizes of hunter-gatherers (levels 4 and 5; regional and ethnolinguistic populations) are among the largest social groups of all mammals and primates. The only other mammals in Figure 5A with similar body sizes and social groups are all marine mammals, including seven species of Delphinidae (*Stenella attenuata; Delphinus delphis; D. capensis; Peponocephala electra; Lissodelphis peronii; L. borealis; and S. longirostris*). Interestingly, the human body size of ∼60 kg place them near the empirical apex of maximum group sizes observed at any body size suggesting the human ability to form large social groups is partially a serendipitous function of having a body size near the midpoint of mammal body sizes, above which maximum group size declines rapidly. We discuss this further below.

**Figure 5.**
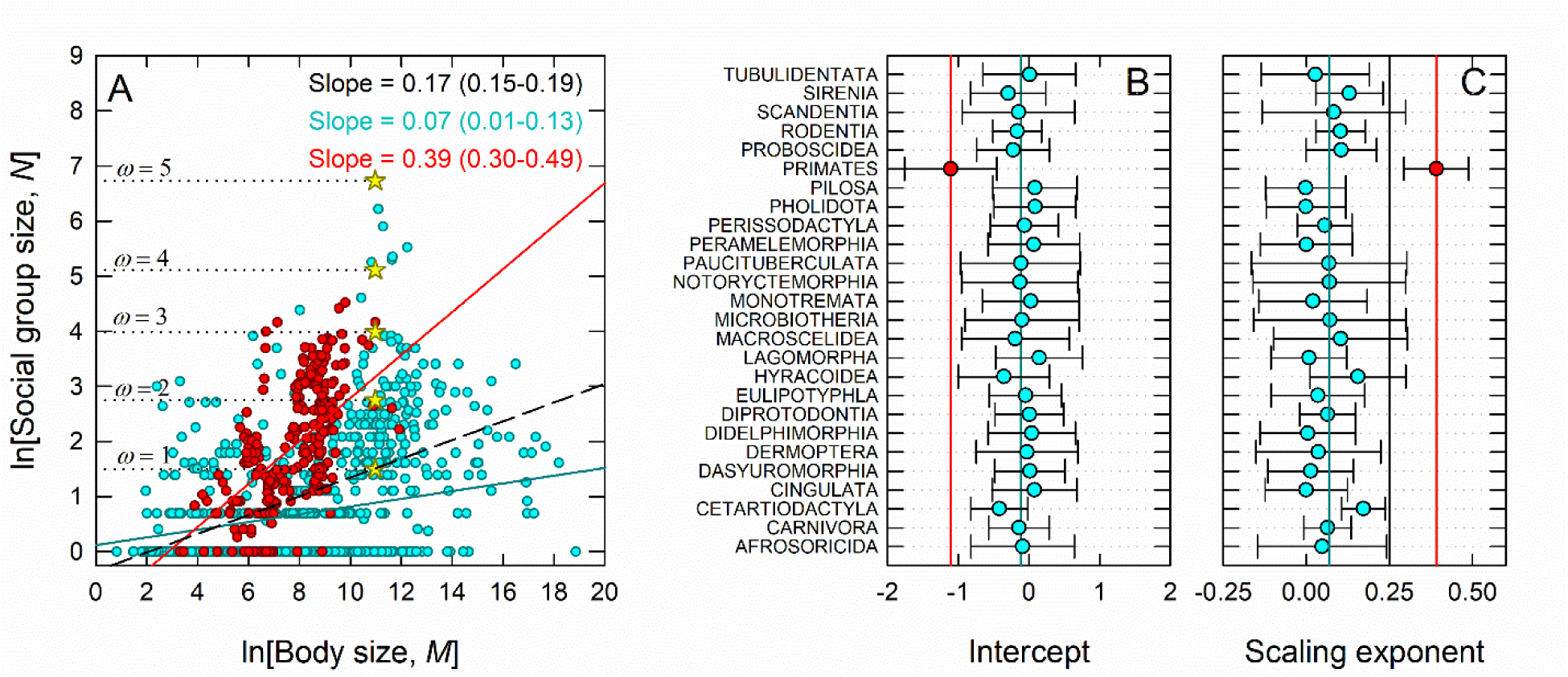
Group size allometries in mammals, primates and humans. A) Group size and body size. B) Intercepts by order from the phylogenetic mixed model. C) Scaling exponents by order from the Bayesian phylogenetic mixed model. The vertical solid black line is the expectation from the null model, equation 2.

### d) Brain size and social networks of brains

A central aspect of group living is cooperation and the increased potential for information sharing among group members. One of the potential cognitive benefits of group living is the ability to interact with more brains. In mammals brain size, *W*, scales with body size as *W* ∝ *M* ^3/4^ (Burger et al., 2019). We see this empirically in figure 6A, where *W* ∝ *M* ^0.73^ (see Table 1). In Figure 6C the BPMM shows that primate brain size (red data point and vertical line) increases with body size significantly faster than other mammals (vertical blue line) (see *ESM* for full results). Therefore, in primates, brains constitute an increasing proportion of body size as body size increases, with the notable exception of Chiropterans (see Smaers et al., 2012). Although the human brain is large for a mammal of 60,000 g at ∼1,350 g, there are several large-bodied mammalian species that have brains absolutely larger than humans, including 26 marine species of Cetartiodactlya (whales and dolphins) and both extant species of Proboscidea; *Loxodonta Africana and Elephas maximus*.

**Figure 6.**
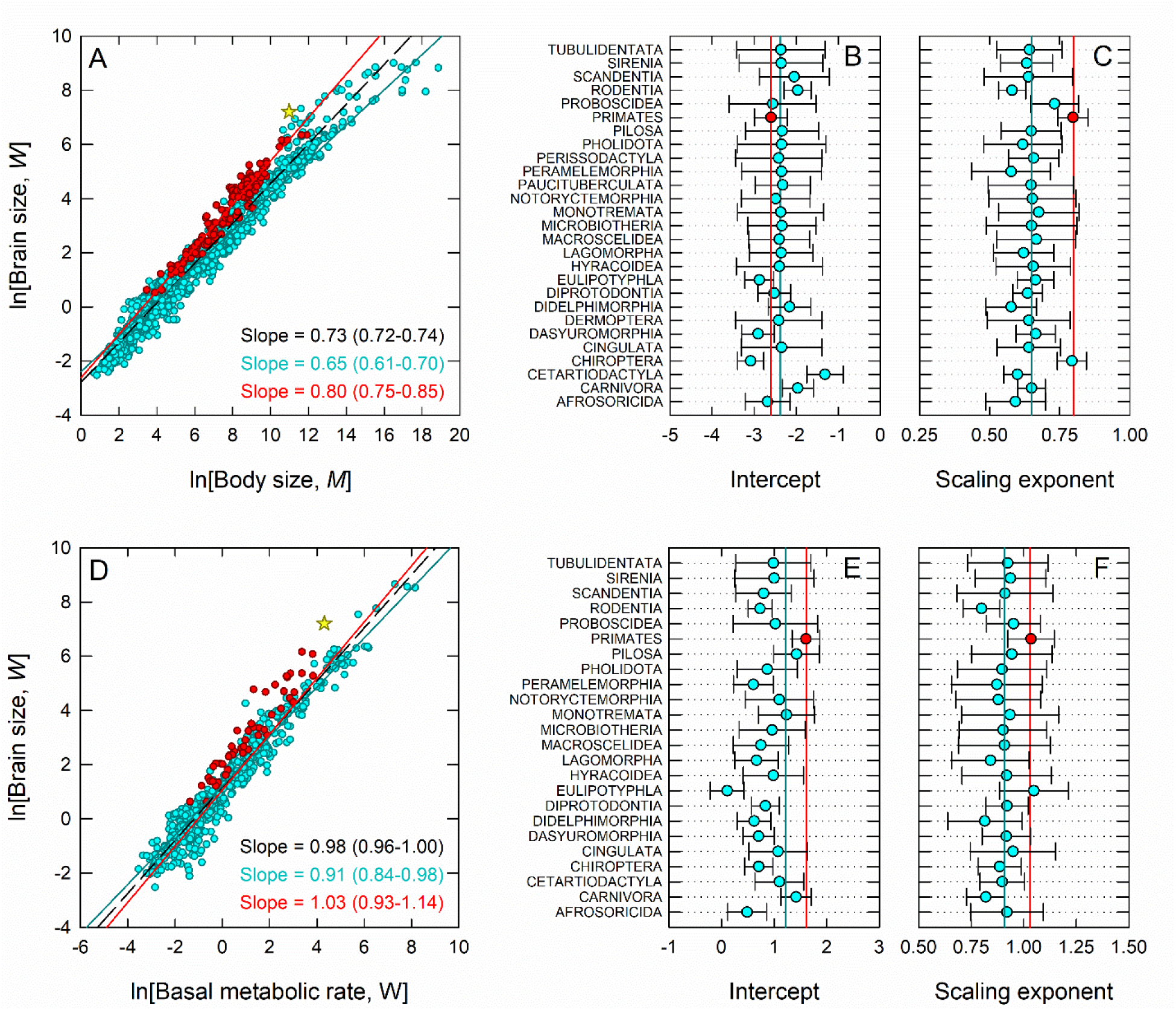
Brain size allometries in mammals, primates and humans. Red data points, lines, and text are non-human primates, blue are non-primate mammals, and yellow diamonds are human. The red and blue lines are the scaling estimates from Bayesian phylogenetic mixed models and the dashed black line and text is the OLS model estimate of the slope. A) Scaling of basal metabolic rate in watts and body size in grams; B) Intercepts from the phylogenetic mixed model by order; C) Scaling exponents from the Bayesian phylogenetic mixed model by order; D-F) Scaling of brain size in grams and body size, and Bayesian phylogenetic mixed model results; G-I) Scaling of brain size and basal metabolic rate, and Bayesian phylogenetic mixed model results.

Consequently mammalian brain size increases linearly with basal metabolic rate, *W* ∝ *B*^1^ as shown in Figure 6D and Table 1; the faster the metabolic rate the larger the brain. Figure 6E shows that primate brains are significantly more expensive to support per gram than other mammal brains, and Figure 6F shows that primate brain size increases significantly faster with basal metabolic rate than other mammals (see ESM for full results). Metabolically expensive primate brains thus become increasingly expensive with size. Figure 6A shows that while human brains are particularly metabolically expensive, so are those of several larger-bodied primates, including *Pongo pygmaeus, Papio Anubis, P. hamadryas, P. papio, Cercocebus torquatus, Erythrocebus patas, and Hylobates lar*.

To consider how brain allometry scales up in social groups we combine the allometric scaling of brain size, *W*, and social group size, *N*, to define the collective brain mass of a social group as *G* = *NW*. An allometric null model for collective brain mass is then

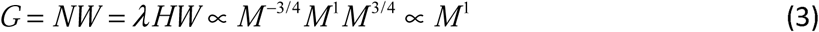

Thus, the null model predicts collective brain mass within species-specific social groups increases linearly with body size. If so, this would have important consequences for group cognition as average brain size increases sublinearly with body size. Therefore, as body size increases brains would be connected within larger social networks, allowing for greater connectivity. However, Figure 7A shows that the scaling of collective brain mass across mammals is sublinear and significantly less than 1 (see Figure 7C), both in the OLS model *G* ∝ *M* ^0.87^ and the BPMM, *G* ∝ *M* ^0.72^. However, for primates the scaling is superlinear, *G* ∝ *M* ^1.20^, and significantly greater than 1 (the vertical black line in Figure 7C) (see *ESM* for full details). Primate collective brain mass increases with body size much faster than other mammals (∼1.7-fold). In addition, primates are the only mammalian order to scale significantly differently than all other mammalian orders (Figure 7C). Hunter-gatherers exhibit among the largest collective brain masses of any mammal or primate (Figure 7A). As with social group size, the only other mammals with similar-sized collective brain masses are species of the Delphinidae family. The phylogeny of collective brain mass across mammals is shown in Figure 8.

**Figure 7.**
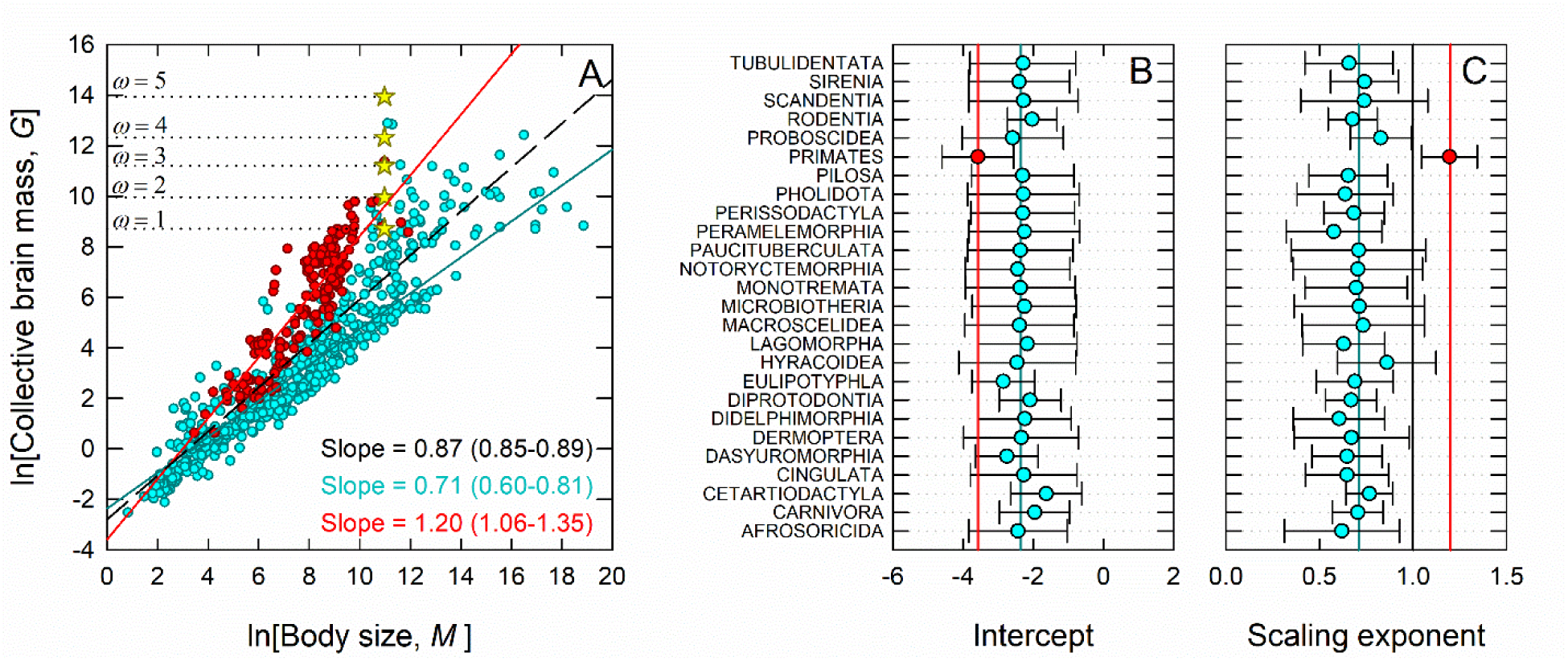
Collective brain size allometries in mammals, primates and humans. A) Collective brain size and body size. B) Intercepts by order from the phylogenetic mixed model. C) Scaling exponents by order from the Bayesian phylogenetic mixed model. The vertical solid black line is the expectation from the null model, equation 3.

**Figure 8.**
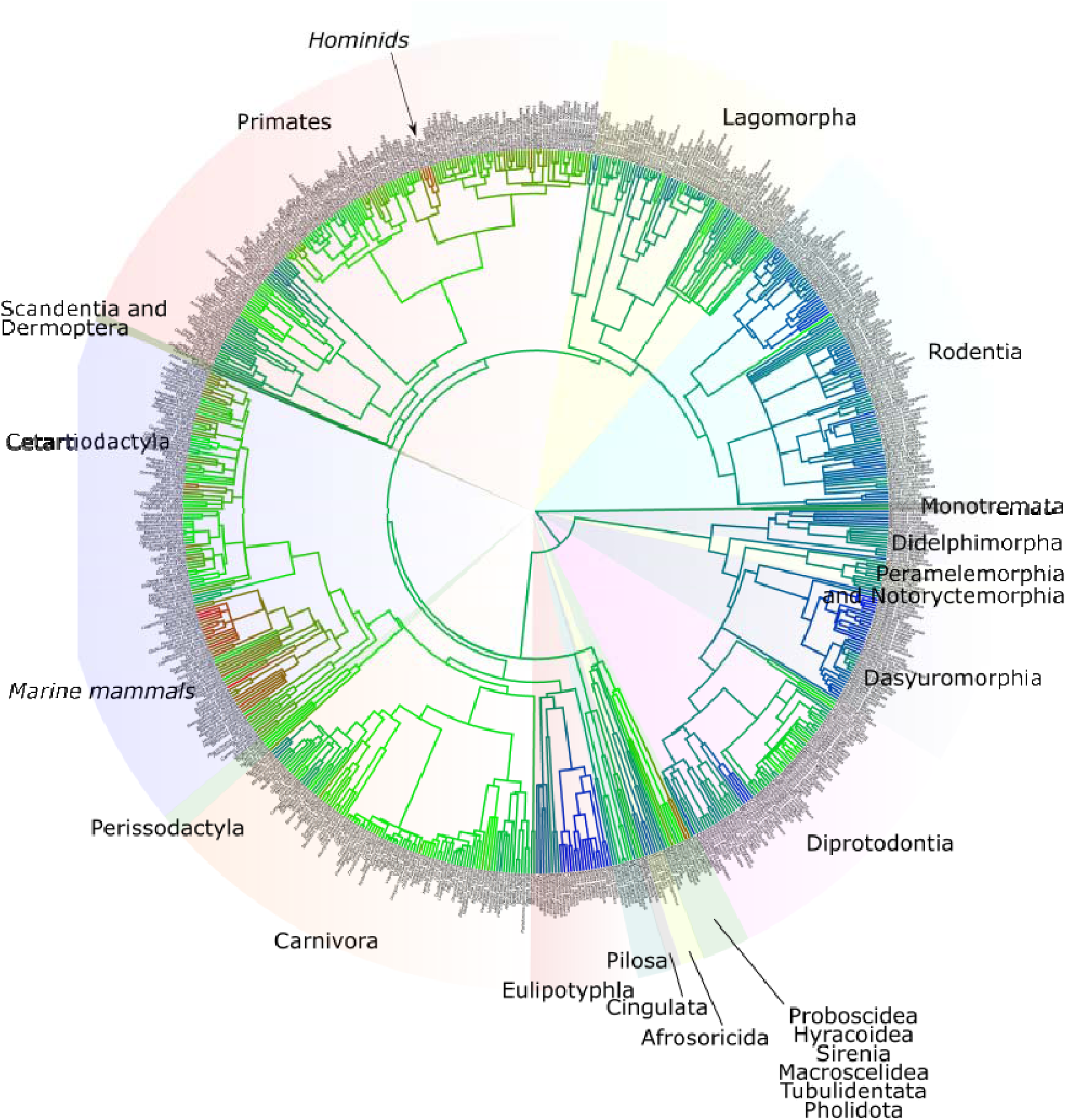
The phylogeny of group brain networks in mammals using continuous character mapping. Colored sectors denote mammal orders.

## 3. Discussion

Primate species form social groups that are among the largest for their body size among mammals. The allometric scaling of primate group size is unique among mammals and the data show that group size increases with body size almost twice as fast than in other mammal (Figures 6A and C). The data show that this occurs because primates support greater densities of individuals than most other mammals (Figures 3A and 4D). The analysis of the null model we derive in this paper from ecological theory shows that primates form groups faster than would be expected from the random encounter rates of individuals within shared home ranges. As species in all other mammal orders tend to form groups slower than the null expectation, the results we report here show a strong statistical signature of a generalized primate prosociality not found in other mammals and which results in the unique allometry of social group scaling in primates.

Hunter-gatherers have a wide range of social group sizes (Figure 5A), including some of the largest observed in all mammals. The most common residential group sizes used during the course of a year are the dispersed and aggregated bands (levels 2 and 3 in Table 2 and Figure 5A) that reflect the seasonal fission-fusion cycle of families as they aggregate into different group sizes over the course of a year (Binford, 2001). Figure 5A shows the sizes of these groups are much as expected for a primate of ∼60 kg. However, because individual hunter-gatherer bands are embedded within larger social networks commonly of hundreds to thousands of people, all of which interact at various rates (Hill et al., 2014), these social group sizes are among the largest observed of any mammal species. While some mammal species aggregate into large collectives of many thousands of individuals, such as herds of artiodactyls or colonies of chiropterans in roosts, these aggregations are not social networks of co-residing individuals, but aggregations brought about by environmental conditions that are temporarily capable of supporting large focal densities of individuals. Analogous aggregations in human societies may be large-scale ritual events held over a number of days, or sporting events that aggregate tens of thousands for short periods of time. Of course, the largest human social groups in the 21^st^ century connect billions of individuals though social media, cell phone networks, religious identities, languages, or the politics of nation states.

The self-organization of hunter-gatherer meta-populations into a fluid arrangement where families aggregate into different sizes of social groups at multiple levels results in a highly flexible social organization. On the one hand, the number of co-residing families in a band is often no more than 3-6 (Hamilton et al., 2007b, 2018), perhaps 10-20 individuals (Hill et al., 2011), a size which may optimize group return rates while minimizing competition for finite resources and the risk of free-riders, for example. On the other hand, by maintaining a fluid structure of fission and fusion among meta-populations composed of hundreds of people with some form of shared identity, information flows at much larger scales, often including multiple neighboring metapopulations (see Bird et al., 2019). Thus individuals living in small social groups on a daily basis receive the long-term benefit of being embedded within a much larger social network.

A consequence of the unique allometric scaling of primate group sizes is the similarly unique scaling of collective brain masses in social groups of primates (Figure 6A). This is because both social group size and brain size scale faster in primates than in all other mammals (Figure 6C) and so collective brain mass in primates diverges from all other mammals as body size increases. Therefore, at all body sizes, primates have both larger brains and larger social networks for that brain to interact with than other mammals. Importantly, the rate of allometric increase of collective brain mass in primates is superlinear (i.e., slope > 1). This means that in primate species increasingly larger brains are embedded within even larger social groups. Because group connectivity increases multiplicatively with size (i.e., *c* ∼ *n*^2^) then larger brains have a multiplicatively greater potential for interaction than smaller brains. Across all other mammal orders the collective brain mass within social groups increases sublinearly with body size (Figures 6A and C), because larger brains occur in social group sizes that increase with body size only very shallowly (if at all) (Figure 6A and C).

Because the number of neurons scales positively with brain size in mammals (Herculano-Houzel, 2012; Herculano-Houzel et al., 2015), a social group with a larger collective brain mass can potentially recruit more brains with more neurons to perform a collective behavior, and so it reasonable to hypothesize that larger social networks will have a greater potential for group cognition and collective computation. If so, primates would be predicted to exhibit a greater potential for group cognition than non-primate mammals. It is perhaps not surprising then that the most evidence we have for advanced group cognition and collective computation in mammals comes from one of the largest-bodied primates.

Intriguingly, Figure 5A suggests that the ability of humans to form large social groups may be partially a function of its position on the mammal body size spectrum. The only other mammals with similar-sized social groups are several species of Delphinidae, all of whom have similar-sized bodies to humans. From the smallest terrestrial mammal, *Suncus etruscus*, the Estruscan shrew at ∼2 g, maximum social group size increases steadily up to the human body size of ∼60,000 g, but thereafter declines rapidly. This suggests that while average group size may well be a function of body size and ecology (as described in our model), as body size increases there are shifting trade-offs in the ability of ecosystems to support the largest aggregations of larger-bodied mammals: above a certain body size (apparently near the midpoint of the body size spectrum) large groups are increasingly difficult to maintain and maximum group sizes decrease monotonically. This may because about half of the mammal species with body sizes larger than humans are marine mammals (data not shown), which have different ecological constraints to maintaining aggregations than terrestrial mammals.

The estimated clade age of primates is on the order of ∼53-88 MYA (Springer et al., 2012; O’Leary et al., 2013), and so the distinctive evolutionary trajectory of primates was a pathway that emerged early in the post-Cretaceous mammalian radiation. The hominin branch emerged ∼6-7 million years ago within the primate tree. Among other traits, the hominin clade saw not only increasing brain size but increasing neuroanatomical specialization (Lieberman, 2011) that ultimately allowed for self-expression (Berwick and Chomsky, 2016; Tattersall, 2019) and the transfer of social information among individuals through gestures, symbols, and language (Tomasello, 2010; Clark and Toribio, 2012; Everett, 2017). Some researchers suggest that modern human-like social complexity may have emerged with the first major expansion in hominin brain size ∼2 MYA with *Homo erectus* (*senso lato*) (Deacon, 1998; Wrangham, 2009; Herculano-Houzel, 2016; Everett, 2017). However, the point at which complex social networks of the form we see in ethnohistoric hunter-gatherers first appear in the archaeological record is unclear.

It has been argued that the complexity, diversity, and specialization of the primate brain evolved in response to the selective pressures of living in complex 3-dimensional, arboreal environments (Passingham and Wise, 2012). Complex ecological environments are also complex social environments as the successful exploitation of ecological niches necessarily involves the successful negotiation of conspecifics (Passingham, 2008; Passingham and Wise, 2012; Wittmann et al., 2018). Recent research in neuroscience suggests that the ability of brains to restrict attention to a subset of stimuli in complex environments resulted in the capacity for self-awareness of mental state, which in turn became awareness of the mental state of others (Frith and Frith, 2006; Graziano, 2017). If this is an accurate mechanistic model, the results we show here suggest that the evolutionary consequences of these dynamics led to feedbacks between group size and brain size that resulted in the unique allometries of group size and collective brain mass in primate species.

## 4. Conclusion

The unique allometric scaling of group sizes in primates leads to the superlinear scaling of collective brain masses. This superlinearity results from the faster allometric scaling of both brain size and group size in primates than other mammals. Thus primates have both larger brains and larger social networks than other mammals of a similar body size. Consequently, as large-bodied primates, human hunter-gatherers have the largest extended brain networks of any mammal. These networks have grown in size, scale, and complexity over human evolutionary history, particularly the last few thousand years to the extent that the majority of the human species is now connected globally by various overlapping social networks. Ultimately, the scale of these networks have deep evolutionary roots in mammalian allometry, primate ecology, and the serendipity of human body size.

## 5. Acknowledgements

We thank Fernando Campos, Briggs Buchanan, Robin Dunbar and various anonymous revierwers for comments on earlier drafts of this manuscript. We also thank Eric Rupley, Sam Scarpino, and Giovanni Petra for extended discussions of many of the topics covered in this paper. This research did not receive any specific grant from funding agencies in the public, commercial, or not-for-profit sectors.

